# Selective changes in cortical cholinergic signaling during learning

**DOI:** 10.1101/2025.08.29.673096

**Authors:** Josue Ortega Caro, Hannah M. Batchelor, Sweyta Lohani, David van Dijk, Jessica A. Cardin

## Abstract

Cognition relies on the function of local and long-range neural circuits in the neocortex, which are dynamically regulated by neuromodulatory signals including acetylcholine (ACh). Recent work suggests that the role of acetylcholine varies across cortical areas and behavioral states. In addition, the precise role of cholinergic signaling during learning and plasticity remains unclear. We performed simultaneous, dual-color mesoscopic imaging of ACh and calcium signaling in the neocortex of awake mice across multiple stages of visual task learning to identify learning-related plasticity in neural activity and cholinergic signaling. Using a novel Pattern Reconstruction and Interpretation with a Structured Multimodal transformer (PRISM_T_) model combining masked autoencoding with multimodal causal attention, we identified key cortical regions that reorganize with learning. We found that cholinergic signaling exhibited spatiotemporally selective plasticity in frontal cortical subnetworks during visual perceptual learning. Overall, our findings demonstrate the utility of transformer models for generating biologically interpretable insight into large multimodal datasets and link long-term functional reorganization of cortical networks to enhanced task performance.

## Introduction

Local and long-range circuit interactions in the neocortex underlie cognition and complex behaviors and are regulated by multiple neuromodulatory systems, including acetylcholine (ACh). Indeed, cholinergic signaling contributes to attention, arousal, sensory processing, and learning and memory ^1-9^. However, the relationship between ACh and cortical activity is both area- and state-dependent^10^, posing a complex challenge to developing a comprehensive model of cholinergic function. The precise role of cholinergic signaling in dynamically reorganizing large-scale networks thus remains poorly understood. Moreover, addressing these questions requires an analytical framework that can integrate information across multiple spatial scales, capture complex temporal dependencies associated with plasticity, and jointly analyze multimodal signals such as neuronal activity and cholinergic dynamics.

A major obstacle to examination of longitudinal changes in large-scale multimodal dynamics has been the development of modeling frameworks capable of extracting meaningful trial-by-trial dynamics from high-dimensional, multimodal recordings. Traditional approaches, including linear regression, generalized linear models, or dimensionality reduction methods such as PCA and factor analysis^11^, have provided important first insights into large-scale cortical activity, but they often assume linearity and lack the capacity to capture nonlinear or context-dependent dependencies. Dynamical systems approaches, including recurrent switching linear dynamical systems (rSLDS)^12^, and latent factor models, such as non-negative matrix factorization (NMF)^13^, have enabled the characterization of low-dimensional trajectories underlying neural population activity^14^. These models have been powerful in uncovering smooth latent representations of neural dynamics during motor control and decision-making tasks^15–22^. However, their sequential processing and strong assumptions about latent structure limit their ability to flexibly capture the heterogeneous and distributed nature of multimodal cortical signals.

More recent deep learning approaches, particularly recurrent neural networks (RNNs) such as gated recurrent units (GRUs) and long short-term memory (LSTMs)^23^, latent factor analysis via dynamical systems (LFADS)^24^, and state-space models (SSMs), have been applied to neural data to model sequential dependencies and predict behavior^25–28^. While these models outperform linear methods, they remain constrained by their reliance on sequential updating, which makes it difficult to capture long-range dependencies and interactions across spatially distant brain regions. In addition, most machine learning approaches generally lack mechanistic interpretability, making it challenging to identify which specific brain regions or modalities drive behavioral predictions.

Transformer architectures offer a compelling solution to these challenges. By employing self-attention mechanisms, transformers can flexibly model global dependencies across space, time, and modality, allowing them to integrate information from widely distributed cortical regions in parallel. Unlike recurrent or state-space models, transformers do not rely on stepwise updates but instead compute relationships between all inputs simultaneously, making them particularly well-suited for the high-dimensional, multimodal, and non-stationary structure of neural data^29^. In addition, the attention mechanism provides a principled and interpretable window into model function^30^, enabling the potential identification of cortical regions and timescales most critical for predicting behavioral outcomes.

Here, we developed the Pattern Reconstruction and Interpretation with a Structured Multimodal transformer (PRISM_T_) model to assess joint mesoscopic recordings of neural dynamics and cholinergic signaling during a visual learning task in mice. By combining masked autoencoding with multimodal causal attention, PRISM_T_ both outperforms existing recurrent and dynamical systems models in reconstructing and predicting neural activity and provides interpretable mapping of key cortical regions that reorganize with learning. We found that cholinergic signaling exhibited spatiotemporally selective plasticity during two stages of visual learning, particularly in frontal cortical subnetworks. Cholinergic signaling patterns became tightly correlated within these subnetworks in associated with enhancement of behavioral performance during visual perceptual learning. Our results highlight the value of transformer-based modeling for both predictive accuracy and biological interpretability, offering a new path forward for the analysis of distributed multimodal brain activity.

## Results

### Multimodal imaging during visual task learning

To simultaneously monitor neuronal activity and cholinergic signaling in the neocortex of awake mice, we expressed the red fluorescent calcium indicator jRCaMP1b^31^ and the green fluorescent ACh sensor ACh3.0^32^ throughout the brain via neonatal injection of AAV vectors into the transverse sinus^33, 34^. This approach results in cortex-wide, uniform expression of both reporters reports neuronal calcium activity and cholinergic signaling^10^. We then performed mesoscopic imaging^35^ of both reporters through the intact skull of mice that were head-fixed and freely running on a wheel (Fig. 1a). We removed hemodynamic and motion-related artifacts using a regression-based approach we previously developed^10^. Adult mice were trained through multiple shaping stages to lick for a water reward when they detected a drifting grating stimulus on a screen (Fig. 1b-c). After initial shaping, mice were trained in Stage 1 to restrict licks to the response period during stimulus presentation and punished with an extension of the inter-trial interval for false alarm licks (FA) outside the response window (Fig. S1a). After behavioral performance reached criterion for Stage 1, mice moved to Stage 2, in which the visual contrast of the grating stimulus varied across trials (Fig. 1b-c, SFig. 1b-c).

**Figure 1:**
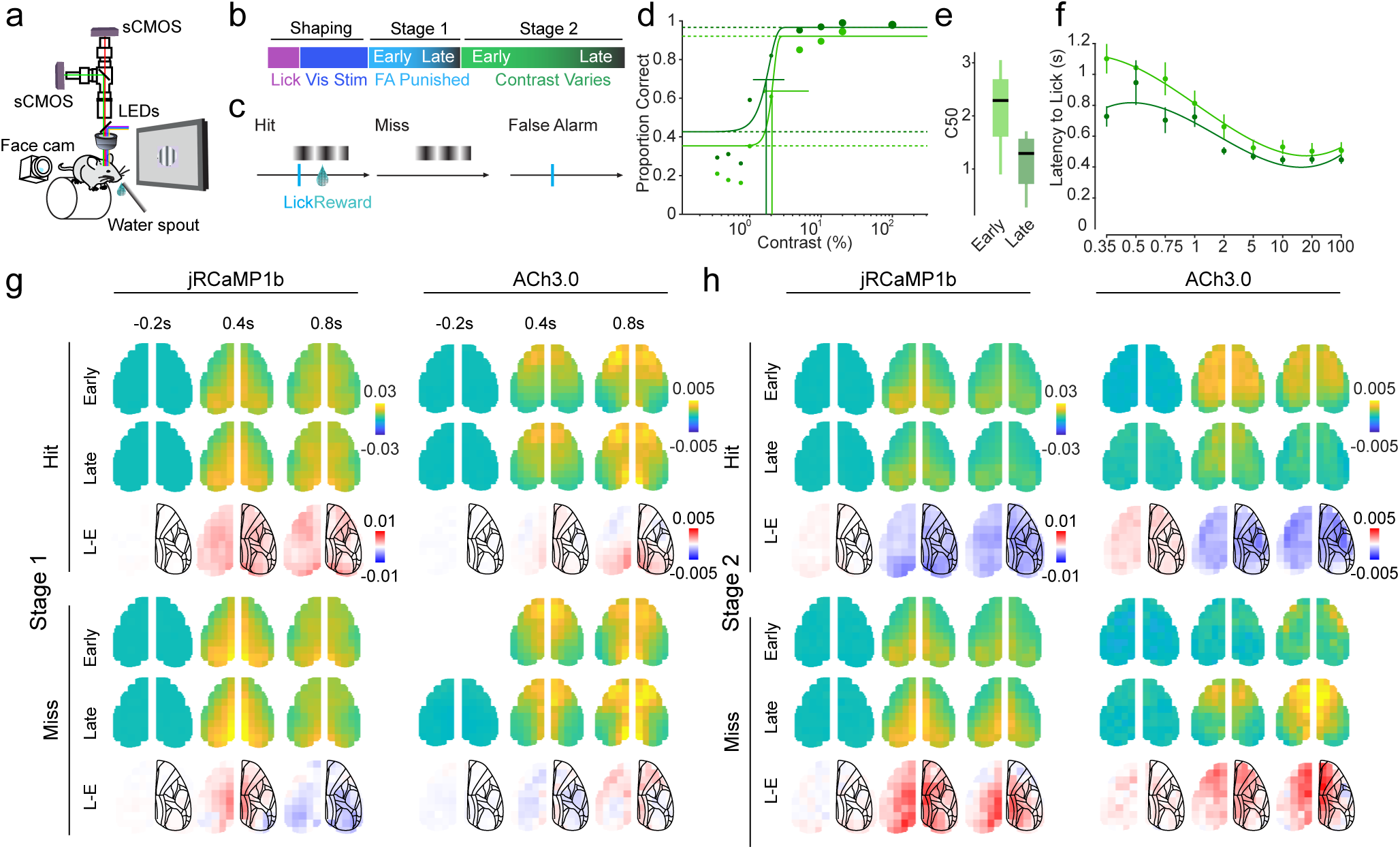
Cortical imaging across multi-stage visual task learning. **a.** Schematic of the dual wavelength, widefield imaging setup with head-fixed mouse on a freely moving running wheel. **b**. Behavioral training included shaping and task learning stages. **c**. The Stage 2 task included three trial types: Hit trials, in which the mouse detected a drifting grating and correctly licked to receive a water reward, Miss trials, in which the mouse did not detect the stimulus and failed to lick, and False Alarms, in which the mouse licked in the absence of a stimulus. **d.** Population average psychophysical performance curves for the first four sessions (Early) and the last four sessions (Late) in Stage 2. Dashed lines denote the false alarm rate and maximum performance and vertical lines denote the contrast at which 50% of trials were correct (C50) for each period. **e**. Population average C50 during Early and Late Stage 2 sessions. (n = 5 mice). **f**. Population average latency to first lick in response to stimuli of each contrast during Early and Late Stage 2 sessions ( n = 5 mice). **g.** Cortical imaging during Stage 1 training for jRCaMP1b (left) and ACh3.0 (right) signals across the dorsal cortex. Upper panels denote Hit trials and lower panels denote Miss trials. In each case, the upper rows shows the average cortical signal at −0.2s, 0.4s, and 0.8s relative to onset of a 100% contrast visual stimulus during Early and Late sessions. The lower row shows the change in cortical activity (Late-Early). Overlays show the Allen Atlas CCv3 anatomical map. **h**. Same as panel g, for ACh3.0.

Consistent with our previous observations^36, 37^, we found that mice exhibited robust perceptual learning during Stage 2. The perceptual threshold, or C50, for visual detection decreased significantly between early (first 4 sessions) and late (last 4 sessions) performance during Stage 2 learning (Fig 1d-e). Across the same time period, mice exhibited a decrease in their latency to first lick response to a visual stimulus (Fig. 1f). Together, these data suggest that mice reliably learn both the Stage 1 and Stage 2 tasks.

In parallel with measuring behavioral outputs, we performed mesoscopic imaging of jRCaMP1b and ACh3.0 signals throughout task learning (Fig. 1g). We observed robust neural responses to task elements in both signals. Initial analysis showed little change in amplitude of either signal in response to the visual stimulus during Stage 1 learning (Fig. 1g) but decreased and increased signals during Hit and Miss trials, respectively, during Stage 2 learning (Fig. 1h). However, the highly dynamic and multimodal nature of cortical activity across multiple timescales poses a challenge for comprehensive analysis.

### Multimodal transformer model captures neural dynamics and trial-to-trial behavior

To address the complexity of the multimodal imaging and behavioral dataset, we developed a model for Pattern Recognition and Interpretation with a Structured Multimodal Transformer (PRISM_T_). Transformers are neural network architectures that excel at handling long-range dependencies through self-attention mechanisms. Unlike recurrent models such as gated recurrent units (GRUs)^38^ or state-space models (SSMs)^23^, which process sequences sequentially, transformers can capture global relationships between data points in parallel, making them well-suited for modeling spatiotemporal neural signals.

We trained PRISM_T_ using a masked autoencoder approach, in which information is removed from the input and predicted from the remaining information. Masked autoencoders are advantageous in learning underlying structures in the data because the model is forced to infer missing components, enhancing its ability to generalize across trials. We used the jRCaMP1b and ACh3.0 signals imaged across the dorsal cortex during stimulus presentation (1 second) as input (Figure 2a). The signals were segmented into a grid consisting of 9×9 pixels per timepoint. We performed 3 different masking strategies: random masking of 90% of patches, future masking, or modality masking (Figure 2b). Performance was computed for new mice that the model did not see during training, highlighting the ability of the model to generalize. PRISM_T_ outperformed recurrent models like GRUs and SSMs in reconstructing missing neural activity in random and future prediction in both Stage 1 and Stage 2 (Figure 2c-d). The model also successfully recovered fine-scale neural activity patterns, even when large portions of the data were masked. Future masking revealed that PRISM_T_ maintained enhanced performance in long-scale temporal prediction and across stimulus contrasts compared to SSM and GRU models (Figure 2e, Figure S2b). Random masking and forward masking were critical for model performance (Figure S2e). Furthermore, the predictability of intertrial intervals was similarly high and reducing the input information (200ms) did not affect performance (Figure S2c-d). Single-trial predictions for jRCaMP1b and ACh3.0 signals showed high temporal fidelity to recorded data (Figure 2f). To examine the effect of mouse task learning on model performance, we separated trials from sessions early in training (first 4 days) and late in training (last 4 days) for Stage 2. We measured the predictability (r2) of neural dynamics between early and late periods during training in each cortical area for the random masking reconstruction. We found that the predictability of neural dynamics increased as the mice became more expert on the Stage 2 task (Figure 2g). Finally, PRISM_T_ outperformed other models in trial-to-trial performance prediction, distinguishing between hit and miss trials across different learning stages and visual contrast levels (Figure 2h, Figure S2a). Performance using JRCaMP1b and ACh3.0 signals together exceeded that using either signal alone (Fig. 2h), suggesting that incorporating multimodal signals improved the model’s ability to predict behavioral outcomes. Together, these results indicate that PRISM_T_ accurately predicts neural and behavioral signals and generalizes well to unfamiliar mice.

**Figure 2:**
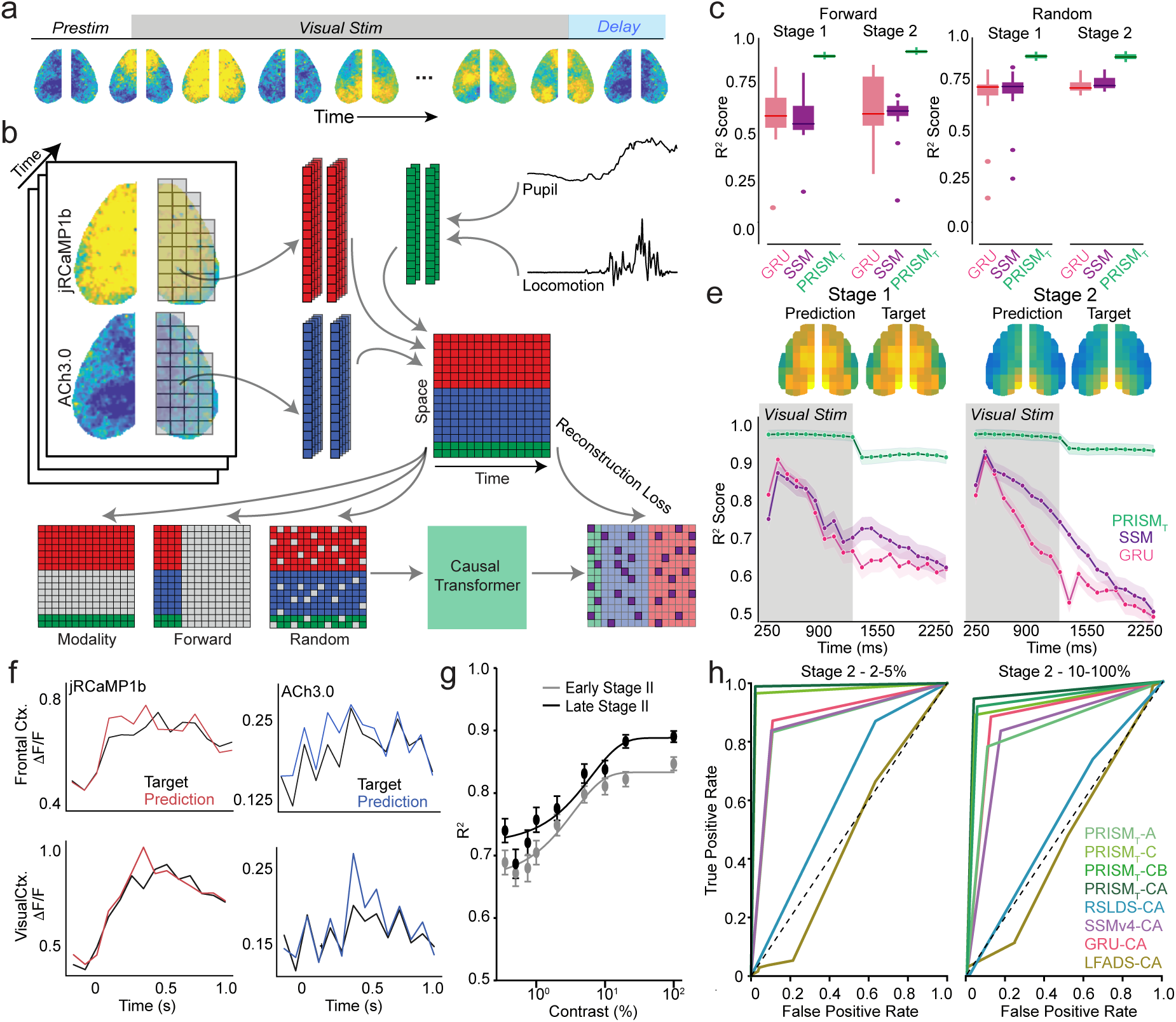
PRISM_T_ predicts neural and behavioral dynamics through learning. **a.** 1 second of widefield calcium and acetylcholine imaging after stimulus onset is input into the model. **b.** The model is pretrained via a Masked Autoencoder. Trials are sampled from 0.2 seconds before stimulus onset to 1.2 seconds after stimulus onset. First jRCaMP1b (Ca), and ACh3.0 (ACh) signals are segmented into non-overlapping patches based on a grid consisting of 9×9 squares (blue and red). Then random segments are masked (grey) and the model needs to reconstruct all unobserved patches. For trial-to-trial performance prediction, the model is finetuned to distinguish between: True Positive vs False Positive or True Positive vs False Negative. **c.** Multimodal Transformer using jRCaMP1b and Ach3.0 data (PRISM_T_) outperforms other multimodal models (GRU, State Space Model (SSM)) on forward masking jRCaMP1b Reconstruction on the validation set mice (N=5). **d.** PRISM_T_ outperforms GRU and SSM in forward prediction on unseen mice (n=80 recording sessions, 3200 trials, 5 mice, 5-fold mice cross validation, p< 0.05, WSR Test). **e.** R2 per timepoint for different models. Models trained on only 1 second are tested on 2 seconds of temporal prediction (n=80 recording sessions, 3200 trials, 5 mice) **f.** Example visualization of jRCaMP1b and ACh3.0 reconstruction from primary visual cortex and frontal cortex using PRISM_T_. **g.** Random masking predictability for PRISM_T_ changes with training. **h.** AUC-ROC curve for trial-to-trial prediction of Hit vs Miss, PRISM_T_ outperform single modality transformers and other multimodality models (SSM, GRU, LFADS) on trial-to-trial behavior prediction in 2-5% contrast and 10-100% contrast trials (n=80 recording sessions, 2911 trials, 5 mice, p < 0.01, DeLong’s test).

### Predictability of neural signals varies with behavioral context

Previous work has shown that the same stimuli presented in different contexts can evoke distinct neural dynamics and may be differently encoded by local and long-range networks^39, 40^. In our training paradigm, mice learned to selectively respond to a single 100% contrast grating stimulus in Stage 1 but also encountered that same stimulus in the context of other stimuli in Stage 2. Using the 100% contrast trials, we examined how behavioral context affected model training and outcomes by taking advantage of Attention Rollout in PRISM_T_. This method computes the relative importance of the different model inputs for the prediction, introducing biological interpretability into a highly complex non-linear model^30^. We found that the attention map used to predict trial-to-trial performance exhibited strong similarity in cortex-wide structure for jRCaMP1b attention across early and late portions of Stages 1 and 2 (Figure 3a). In contrast, we found that the attention map for ACh3.0 changed across learning stages, with increasing attention to frontal cortical areas over time. We further quantified this change in attention by training a new model to predict from which training stage individual 100% contrast trials originated. Whereas the attention map for jRCaMP1b was broadly distributed across cortical areas, the map for ACh3.0 focused attention in primary and secondary motor cortex (Figure 3b), suggesting broadly distributed changes in calcium dynamics but locally specific changes in cholinergic activity across learning. To test whether the jRCaMP1b and Ach3.0 dynamics that predict trial-to-trial behavior change across learning stages, we trained 4 models for each period and tested them on the other periods. All models had above-chance performance for trial-to-trial prediction in the other training periods (Figure 3c), indicating that similar dynamics are associated with trial-to-trial behavior across training stages for 100% contrast trials. Finally, we quantified the change in predictability of neural dynamics within each training period. We found broadly distributed increased predictability for both jRCaMP1b and ACh3.0 dynamics across mouse learning in Stage 1, but a selective increase in predictability for ACh3.0 dynamics in frontal cortical areas during perceptual learning in Stage 2 (Figure 3d). Together, these results suggest that spatiotemporal changes in cholinergic signaling across the cortex may play a key role in learning-related changes in behavioral output.

**Figure 3.**
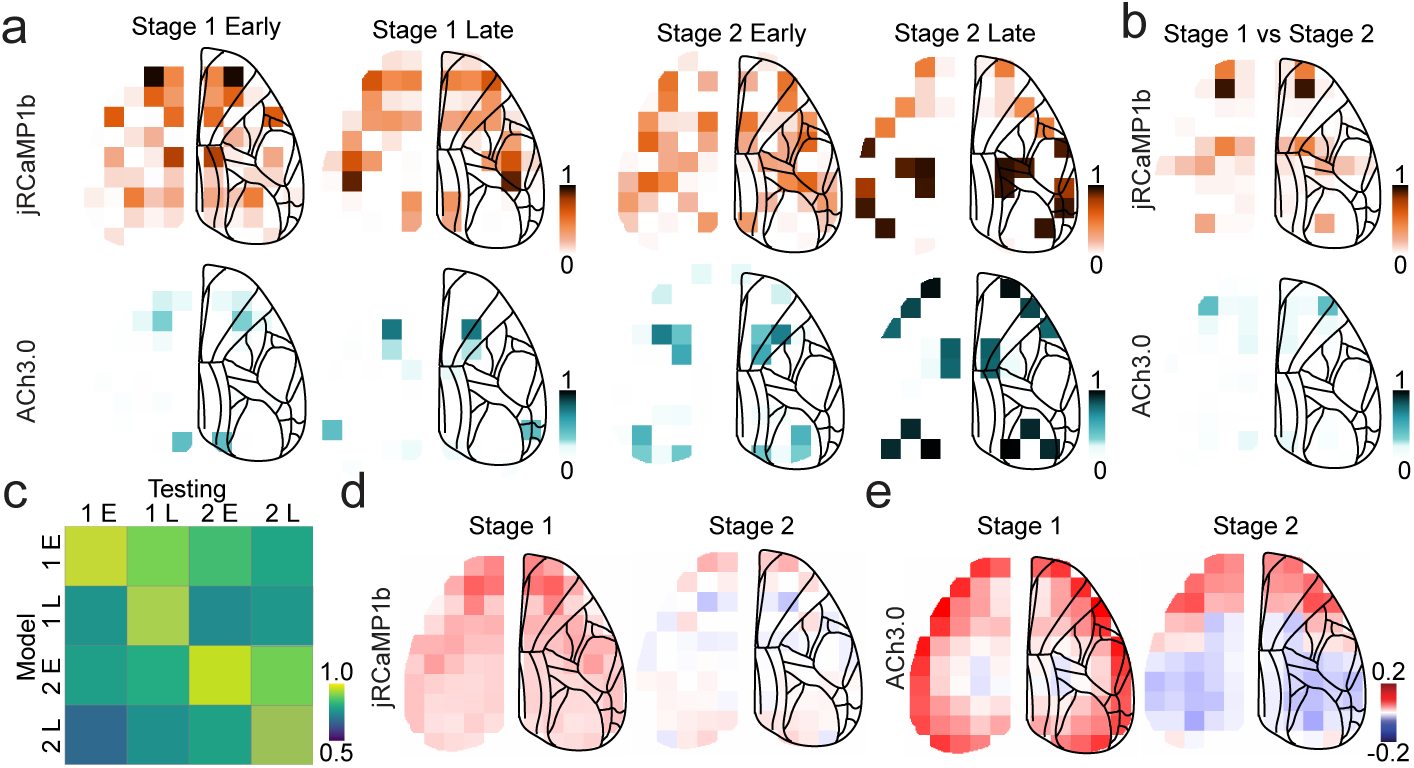
Context dependent dynamics during 100% contrast trials. **a.** Self-attention values for prediction of Hit vs Miss trials using jRCaMP1b (orange) and ACh3.0 (cyan) during stimulus presentation for Early Stage 1 (Day 1-4), Late Stage 1 (last 4 days), Early Stage 2 (Days 1-4), and Late Stage 2 (last 4 days) for 100% contrast trials (per parcel n=32 recording sessions, 1000 trials, 5 mice, p < 0.05, permutation test). **b.** Self-attention values for predicting if a trial comes from Late Stage 1 or Late Stage 2 using jRCaMP1b (orange) and ACh3.0 (cyan) during stimulus presentation for 100% contrast trials (per parcel n=22 recording sessions, 550 trials, 5 mice, p < 0.05, permutation test). **c.** Trial-to-trial prediction across stages. Vertical names indicate the stage the model was trained on, horizontal indicates the stage the model was tested on. Models were able to perform across training stages with above chance performance (5-fold cross validation across n = 5 mice). **d.** Changes in predictability per area between early and late Stage 1 and Stage 2 for jRCaMP1b, respectively. **e.** Changes in predictability per area between early and late Stage 1 and Stage 2 for ACh3.0, respectively.

### Baseline cortical network dynamics predict performance

Previous work has suggested that arousal state, mediated in part by neuromodulatory signals including acetylcholine, robustly regulates cortical network state^41–43^ and task performance^44, 45^.The arousal-dependent baseline state of the cortex prior to each trial (Figure 4a) could thus drive behavioral outcomes and could be subject to plasticity across learning. To examine the role of baseline state on trial outcomes, we tested the predictive power of cholinergic signaling and cortical activity before stimulus onset (Figure 4b). We trained multiple iterations of PRISM_T_ with different inputs and evaluated their performance on trial-to-trial behavioral prediction. Pre-stimulus activity in both the jRCaMP1b and ACh3.0 signals contributed significantly to predicting trial performance. ACh3.0 signals had higher trial-to-trial behavior prediction prior to stimulus onset, whereas jRCaMP1b had higher performance after stimulus onset. To examine how jRCaMP1b and ACh3.0 signals in specific cortical regions during the pre-stimulus period contributed to trial-to-trial behavioral prediction, we computed attention maps for near-threshold (2-5%), mid-range (10-20%), and high (100%) contrast stimulus trials in each training stage. The distribution of attention varied between different stages and stimulus contrasts (Figure 4c), suggesting that the impact of pre-stimulus baseline cortical state varies with behavioral context. However, we found only small changes in predictability (r2) for pre-stimulus baseline activity in either the jRCaMP1b or ACh3.0 signal across training (Figure 4d), suggesting that the dynamic structure of the baseline cortical state does not change with learning. To further estimate the stability of network interactions across learning stages, we calculated a correlation-based functional connectivity measure between pairs of cortical areas during the pre-stimulus period. Functional connectivity across the cortex was largely unchanged for pre-stimulus jRCaMP1b and ACh3.0 signals, suggesting little learning-related change to underlying baseline cortical network interactions (Figure 4e, Figure S4b). The role of pre-stimulus cortical state in trial-by-trial behavioral outcome is thus context-dependent but static across task learning.

**Figure 4.**
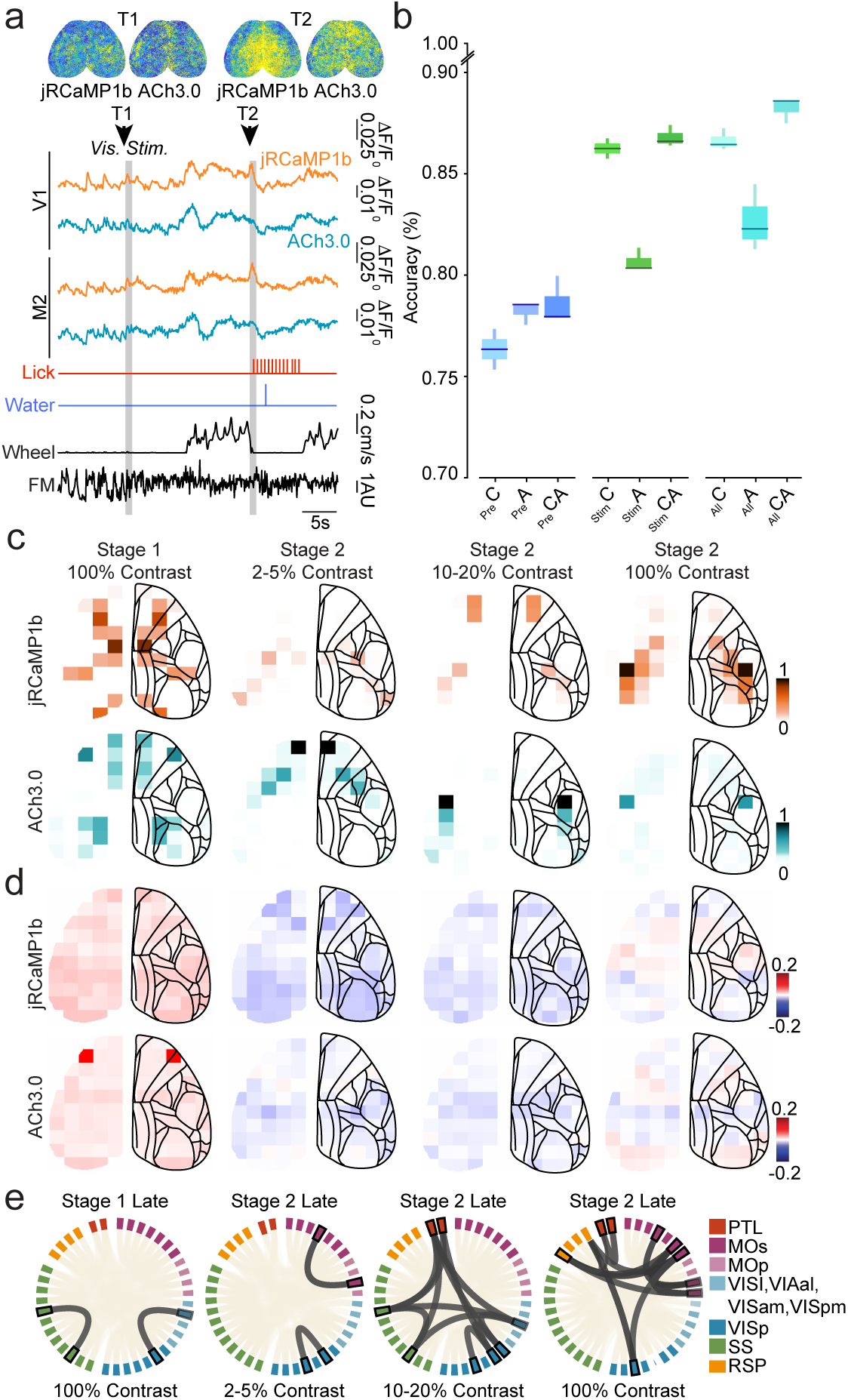
Pre stimulus onset dynamics predicts trial-to-trial mice performance. **a**. Example jRCaMP1b and ACh3.0 traces from pre-trial baseline and trial periods for primary visual cortex and secondary motor cortex. Lower traces show lick, water reward, locomotion on the wheel and facial motion energy. **b.** Prediction of Hit vs Miss Trials using jRCaMP1b (orange) and ACh3.0 (cyan) before stimulus onset for near threshold (2-5%) trials (n=80 sessions, 3200 trials, 5 mice, 5-fold cross validation). **c.** Self-attention values for prediction of Hit vs Miss trials using jRCaMP1b (orange) and ACh3.0 (cyan) during stimulus presentation for late stage 1 (Last 4 days) for 100% contrast trials, late stage 2 (last 4 days) for 2-5% contrast trials, 10-20% contrast trials, and 100% contrast trials (per parcel n=22 recording sessions, 550 trials, 5 mice, p < 0.05, permutation test). Shows qualitative differences in attention across stages and contrast. **d.** Changes in forward masking predictability (Late days minus Early Days) for pre-trial dynamics (500 ms before stim onset) by area for jRCaMP1b and ACh3.0. Red indicates increases in predictability and blue indicates decreases. **e.** Chord plot visualizing statistically significant functional connectivity between model-selected grids (n = 8405, p < 0.05, linear mixture effects model) for Late Stage 1 (100% contrast), Late Stage 2 (2-5% contrast), Late Stage 2 (10-20% contrast), and Late Stage 2 (100% Contrast) for jRCaMP1b signal. Bold indicates model-selected grids, denoted by Allen Atlas CCv3 labels.

### Cholinergic signaling exhibits selective plasticity associated with visual learning

Sensory cue-evoked cholinergic signaling in sensory and frontal cortical areas has been linked to learning of task contingencies and reward prediction^46–49^. Neuromodulatory signals evoked by task events may therefore be targets of learning-related plasticity. However, it remains unclear how such changes might be coordinated across long-range cortical networks. To more closely examine changes associated with perceptual learning, we separated the Stage 2 task trials into near-threshold (2-5% contrast) and mid-range (10-20%) trials and generated attention maps for jRCaMP1b and ACh3.0 signals across early and late Stage 2 sessions. For near-threshold trials, the jRCaMP1b attention distribution during early and late Stage 2 learning were similar, focusing on frontal, somatosensory, and visual cortical areas in both beginner and expert mice (Figure 5b-c). In contrast, the ACh3.0 attention pattern shifted from visual cortex to frontal cortex over this same learning period. Furthermore, attention for both signals shifted towards frontal cortex across learning for mid-range contrast trials. This pattern of attentional shift was robust and emerged from both population and individual mouse analysis (Figure S5a).

**Figure 5.**
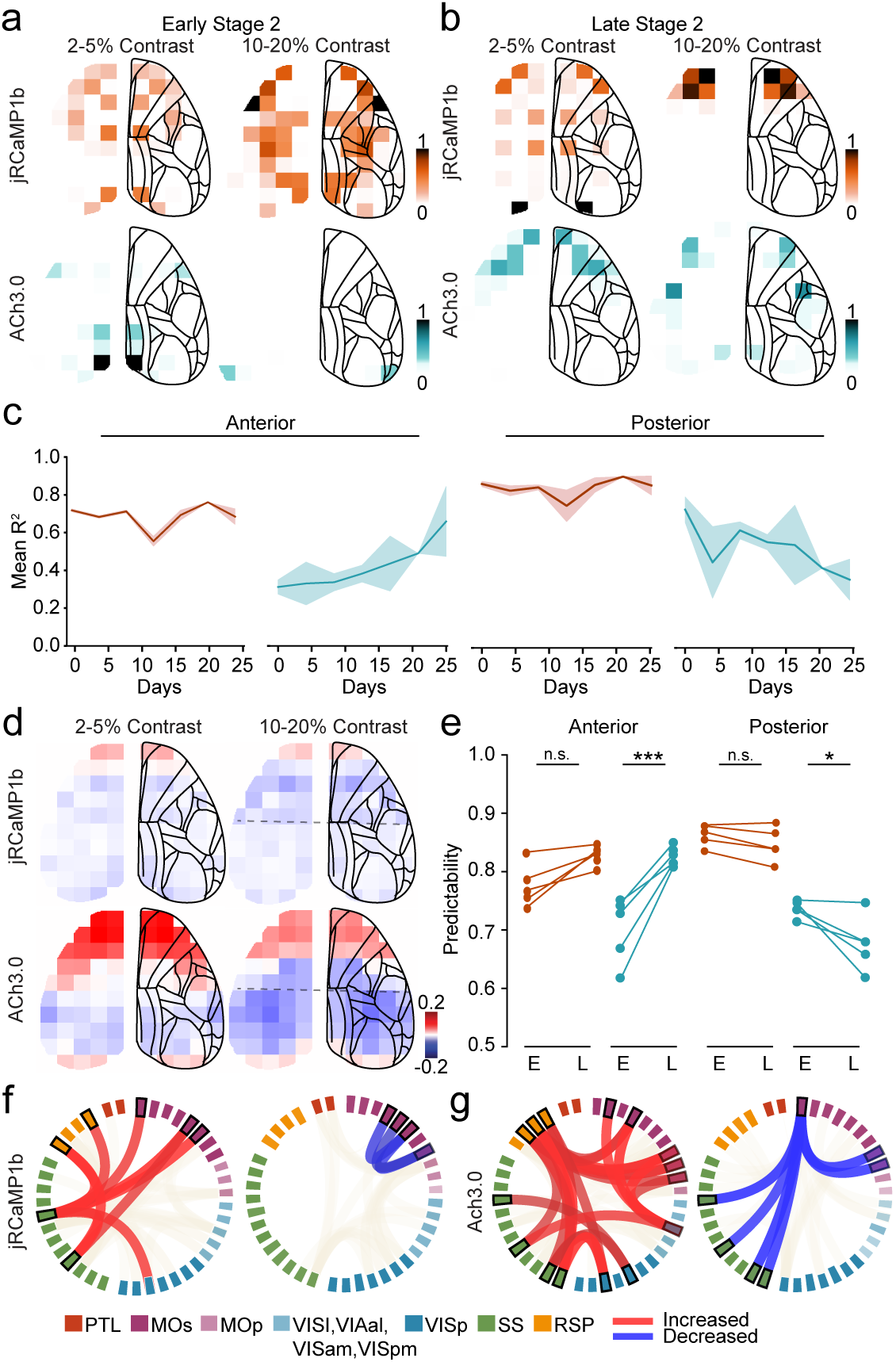
Changes in attention and dynamics associated with learning. **a.** Self-attention values for prediction of Hit vs Miss trials using jRCaMP1b (orange) and ACh3.0 (cyan) during stimulus presentation for Early Stage 2 training days (Day 1-4) for 2-5% and 10-20% contrast trials (per parcel n=20 recording sessions, 802 trials, 5 mice, p < 0.05, permutation test). **b**. Self-attention values for prediction of Hit vs Miss trials using jRCaMP1b (orange) and ACh3.0 (cyan) during stimulus presentation for Late Stage 2 training days (Last 4 days) for 2-5% and 10-20% contrast trials (per parcel n=40 recording sessions, 1043 trials, 5 mice, p < 0.05, permutation test). **c.** R2 across days for random masking during learning between anterior and posterior brain areas for 2-5% contrast trials. Split was selected based by using retrosplenial cortex as a boundary. jRCaMP1b R2 maintains predictability across days for anterior and posterior areas, however, ACh3.0 increases in R2 by days for anterior and decreases for posterior areas. **d.** Changes in forward masking predictability (Late days minus Early Days, 400 ms input) by area for jRCaMP1b and ACh3.0 signals. Red indicates increase in predictability and blue decrease. **e.** Quantification of change in predictability between Early and Late Stage 2. Anterior ACh3.0 (p < 0.001, K-S test), Posterior ACh3.0 (p < 0.05, K-S Test). **f.** *Left*: Chord plot visualizing statistically significant positive changes in functional connectivity between model selected grids (n = 8405, p< 0.05, linear mixture effect model) between Early and Late Stage 2 for 2-5% contrast trials for jRCaMP1b. Bold indicates model-selected grids. Red indicates model grid connections. Beige indicates significant non-model selected grid connections. *Right*: Chord plot visualizing negative changes in functional connectivity between model selected grids. Blue indicates model-selected grids for calcium signal. **g.** *Left*: Chord plot visualizing statistically significant positive changes in functional connectivity between model selected grids (n = 8405, p< 0.05, linear mixture effect model) between Early and Late Stage 2 for 2-5% contrast trials for ACh3.0 signal. Bold indicates model-selected grids. Red indicates model grid connections. Beige indicates significant non-model selected grid connections. *Right*: Chord plot visualizing negative changes in functional connectivity between model selected grids for ACh3.0 signal. Blue indicates model-selected grids.

To relate these changes in model attention to neural dynamics, we again measured the changes in predictability (r2) between early and late Stage 2 periods during training for each cortical area using random masking reconstruction. Based on the spatial shift from posterior to anterior attention, we separated the cortical areas into anterior and posterior groups and quantified the predictability of each signal across learning. ACh3.0 dynamics exhibited increased and decreased predictability in anterior and posterior cortex, respectively, between early and late Stage 2 learning (Figure 5d). In contrast, jRCaMP1b dynamics exhibited little change in predictability across learning, suggesting a selective enhancement of cholinergic signaling patterns.

We further computed the predictability of both signals using forward prediction. Similar to random masking, forward prediction of ACh3.0 signals exhibited increased predictability for anterior cortical areas whereas jRCaMP1b showed no significant change in predictability across learning (Figure 5e-f). Together, these results suggest that the trial-to-trial behavioral outcome prediction, quantified using attention, and trial-to-trial prediction of neural and neuromodulatory dynamics, quantified using predictability, reflect the same underlying changes in cholinergic dynamics due to their spatiotemporal coherence across learning.

To better quantify the changes in coordination of cholinergic signaling across learning that were highlighted by PRISM_T_, we assessed the functional connectivity for all cortical areas across early and late Stage 2 learning. In each learning stage, we compared areas selected for attention by the model to those not selected. After mice reached expert performance in late Stage 2, model-selected areas exhibited enhanced functional connectivity with each other relative to non-selected areas. This enhancement in correlated signaling across areas was significant for cholinergic dynamics in near threshold and high contrast trials but not for calcium dynamics (Fig. 5g-I, Fig. S5d-e). Cholinergic dynamics in the model-selected network overall exhibited increased functional connectivity between retrosplenial, somatosensory, and secondary motor cortex, as well as between primary visual and somatosensory cortex. We further found decreased correlations between motor and somatosensory cortical areas.

## Discussion

Our results demonstrate that PRISM_T_ can capture distributed cortical and neuromodulatory dynamics during learning with both accuracy and interpretability. Benchmarking against established approaches showed that PRISM_T_ more effectively captured trial-level variability and can predict neural dynamics further in time. Beyond performance, the model exposed a clear dissociation between excitatory and cholinergic populations. Context-dependent modulation was stable in neural activity but reorganized markedly in the cholinergic signals, particularly in frontal regions, where they became increasingly predictive of behavior. However, we found that PRISM_T_ trained on the different stages can maintain performance in other periods, indicating that the neural activity that drives trial-to-trial performance for 100% contrast stimulus is maintained throughout. Overall, these results suggest a context-independent role of neural dynamics, in contrast to the context-dependent role of acetylcholine^39, 40^.

In support of the differentiate role of each signal, pre-stimulus dynamics showed a dissociation between learning stages. Neural and neuromodulatory dynamics reorganized broadly during sensory stages. Furthermore, we found that cholinergic signals have higher predictability for trial-to-trial behavior prediction compared to neural signals. This further supports the idea that arousal state, mediated by neuromodulatory signals, regulates task performance^44, 45^.

This differentiable effect of ACh compared to neural activity is the most present in learning-related changes in near threshold trials. Here, we find significant reorganization of acetylcholine dynamics, particularly from posterior to anterior regions, where they became increasingly predictive of behavior. In addition, functional connectivity analyses showed that ACh subnetworks linking retrosplenial, somatosensory, motor, and visual cortices strengthened across training, while excitatory organization remained largely stable. Together, these results highlight cholinergic plasticity as a key mechanism through which distributed cortical circuits adapt during learning.

In sum, our study identifies distinct roles for excitatory and cholinergic populations in shaping cortical dynamics: neural activity maintains distributed scaffolds, whereas cholinergic signals reorganize to modulate behavior via brain state, and flexibly couple cortical subnetworks during learning. By applying PRISM_T_ to multimodal neural recordings, we were able to isolate these processes. This approach provides a generalizable method for dissecting multimodal contributions to circuit dynamics and establishes a mechanistic account of how different signals determine adaptive behavior.

## Methods

### Animals

Male and female wild-type mice (3-6 months) were kept on a 12-h light/dark cycle, provided with food and water ad libitum and housed individually following headpost implantations. Imaging experiments were performed during the light phase of the cycle. All animal handling and experiments were performed according to the ethical guidelines of the Institutional Animal Care and Use Committee of the Yale University School of Medicine.

### Surgical procedures

For widefield imaging, P0/P1 mice were injected in the transverse sinus with fluorescent indicators as described in Lohani, Moberly et al., 2022. Each pup was injected with a mixture of 4 ul of AAV9-syn-jRCamp1b and 4 ul of AAV9-syn-Ach3.0, with half of the total volume injected per hemisphere.

All surgical implantation procedures were performed on adult mice (>P50). Mice were anesthetized using 1–2% isoflurane and maintained at 37 °C for the duration of the surgery. The skin and fascia above the skull were removed from the nasal bone to the posterior of the intra-parietal bone and laterally between the temporal muscles. The surface of the skull was thoroughly cleaned with saline, and the edges of the incision were secured to the skull with Vetbond. A custom titanium headpost was secured to the posterior of the nasal bone with trans-parent dental cement (Metabond, Parkell), and a thin layer of dental cement was applied to the entire dorsal surface of the skull. Next, a layer of cyanoacrylate (Maxi-Cure, Bob Smith Industries) was used to cover the skull and left to cure approximately 30 min at 22–24C room temperature to provide a smooth surface for transcranial imaging. Mice were allowed to recover for 3-5 days after surgery before commencing handling and wheel training.

### Widefield imaging

Imaging was performed as described in Lohani and Moberly et al., 2022. Widefield imaging experiments were conducted ∼ 2 weeks after headpost implantation. Widefield calcium and cholinergic imaging was performed using a Zeiss Axiozoom with a PlanNeoFluar Z ×1, 0.25 numerical aperture objective with a 56-mm working distance. Epifluorescent excitation was provided by an LED bank (Spectra X Light Engine, Lumencor) using three output wavelengths: 395/25 nm, 470/24 nm and 575/25 nm. Emitted light passed through a dual-camera image splitter (TwinCam, Cairn Research) and then through either a 525/50-nm or 630/75-nm emission filter (Chroma) before it reached two scientific CMOS (sCMOS) cameras (Orca-Flash V3, Hamamatsu). Images were acquired at 512 × 512 resolution after 4× pixel binning, and each channel was acquired at 10 Hz with 20-ms exposure. Images were saved to a solid-state drive using HCImage software version 4.5.1.3 (Hamamatsu).

### Behavior Task

Mice were trained to perform a visual detection task with delay, in which visual stimulus and reward delivery were separated in time via a delay to isolate neural and acetylcholine activity corresponding to distinct visual and reward aspects of the task. To motivate mice to perform for water rewards, mice were water restricted to ∼82-88% of their initial body weight. Mice were head-fixed on a wheel and trained to lick at a waterspout after detecting visual stimuli (full screen, temporal frequency: 2Hz; spatial frequency: 0.05 cycles/degree, on a gamma corrected screen with mean luminance ∼ 60 lux) of different visual intensities. Visual stimuli were presented for 1s, which was followed by a delay and a 1s response window. If the mouse licked the spout during this window (hit or correct trial), it received a water drop (reward, 2-3 ul) at the end of the window. If the mouse did not lick during the response window (miss or incorrect trial), reward was withheld. Correct trials were followed by a 3.5s consumption delay and incorrect trials were followed by a 1s delay. Inter-trial intervals (ITIs) were drawn from an exponential distribution to get flat hazard rate and ranged between 1.5 to 10s for stages 1,2,3 and 1.5 to 30s for stages 4,5,6,7,8. ITI licks were punished by ITI timeout, which resulted in the resetting of ITI and prolonging the time to next trial.

Training progressed through several stages that gradually introduced increasingly difficult aspects of the task. In a first shaping stage, mice trained to lick at a waterspout without any visual cue until reaching criterion of >100 licks in a 20m session (∼ 1-2 days).

In the second shaping stage, mice were conditioned to associate visual stimulus (100% visual contrast drifting grating) with reward by always pairing a water drop at the end of the 1s of visual stimulus. The response window started at the offset of visual stimulus and if mice licked during this time, a second water drop was provided. Next, mice were trained solely on an operant task in which reward was contingent on mouse licking during the response window after the visual stimulus. In this stage, there was no delay between stimulus offset and response window. Animals were trained until reaching criterion of >90% correct in a 25m session (∼2-4 days).

Next, a 0.5 s delay was added between stimulus offset and response window. Criterion was met when animals performed with >90% accuracy in a 35m session (∼4-5 days).

In Stage 1 of the final training phase, an ITI timeout as punishment for false alarm licks was introduced. Animals were trained until reaching stable performance of 10-20% false alarm rate in a 50m session (∼5-15 days).

Finally, in Stage 2, a range of low to high contrast visual stimuli (range from 0.35 % to 100%) were presented to get psychometric performance. High contrast trials were oversampled (70% of trials comprised of contrasts over 1.5%) to maintain motivation in the task.

In the non-psychometric stages (1-5), only 100% high contrast visual stimuli were presented. Some mice were removed from further behavioral training if they didn’t meet the performance criteria. Mice were run on the task semi -daily (skipped weekends and holidays), and each session was run for a fixed amount of time (25 min for stage 1, 35 min for stages 2 and 3, 50 min for stages 4 and 5, and 60-75 min for stages 6,7,8), but the session was terminated if the mouse was satiated early.

All behavioral sessions with hit rate below 10% for 100% contrast were removed from analyses. Psychometric sessions were removed if there were fewer than 50 total trials or if the ITI false alarm rate (or hit rate at 0% contrast or 0.35% contrast) exceeded 50% and/or if hit rate for 100% contrast was below 70%.

### Non-instructed Behavior

Non-instructed behavioral variables including locomotion and facial motion were simultaneously measured. A magnetic angle sensor (Digikey) attached to the wheel continuously monitored wheel motion. During widefield imaging sessions, the face (including the pupil and whiskers) was illuminated with an infrared LED bank and imaged with a miniature CMOS camera (Blackfly s-USB3, Flir) with a frame rate of 10 Hz.

### Causal Transformer Architecture

We employed a decoder-only Transformer architecture. The input data consisted of simultaneous calcium and acetylcholine recordings from 41 brain regions, resulting in 82 channels per timestep. Data were segmented into 10 consecutive timepoints sampled at 10 Hz, producing input matrices X in *R*^82*x*10^.

Each scalar value *X*_*i*,*t*_ denotes the signal from channel i at time t. These values were projected into a shared latent space via a learned linear mapping *W*_*e*_ *in R*^1*x*512^:

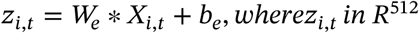

The resulting token embeddings were flattened into a *sequence z*_1_,…, *z*_820_, where 820 = 82 channels * 10 timepoints.

To encode spatial and temporal identity, we added learned positional embeddings *p*_*j*_ in *R*^512^ to each token:

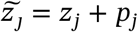

A learned classification token *z*_*CLS*_ in *R*^512^ was prepended to the sequence, forming the input:

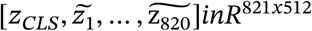

This sequence was passed through L Transformer decoder blocks, each comprising three components:

#### 1. Causal Multi-head Self-Attention

For a hidden state 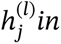 *R*^512^ at layer l, the model computes:

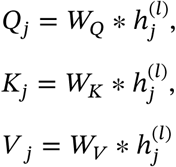

Attention is computed with a causal (lower-triangular) mask:

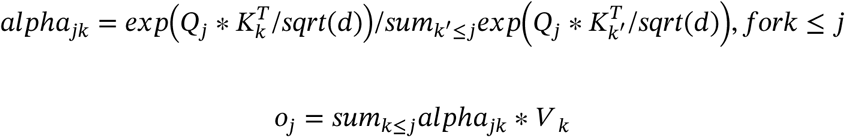

The causal mask follows a step-function pattern: each token at time t attends to all other tokens at time t (i.e., all 82 channels), as well as to all tokens at preceding timepoints t’ < t. This structure preserves spatial dependencies while enforcing temporal causality.

Outputs from all attention heads are concatenated and passed through a linear projection.

#### 2. Feedforward Network

Each token is passed through a two-layer MLP:

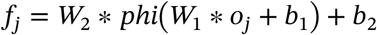

#### 3. Residual Connections and Normalization

Residual connections and layer normalization are applied:

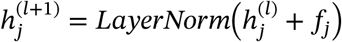

After L layers, the output corresponding to the classification token 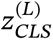 was used as a sequence-level representation.

To facilitate self-supervised learning, we applied masked token modeling: 90% of the non-CLS tokens \*tildez*_*j*_ were replaced with a learned [MASK] embedding. The model was trained to reconstruct the original token embeddings using only the visible context under causal constraints.

This architecture enables autoregressive modeling of distributed neural activity across time and modality, supporting both classification and reconstruction objectives within a unified Transformer framework.

### Training Procedure

The masked autoencoder was trained using the AdamW optimizer, with default hyperparameters η_1_ = 0.9, η_2_ = 0.999, and ɛ = 10^−8^. The loss function was computed only over the masked tokens, encouraging the model to reconstruct unseen inputs based on the visible context. Let *M* ⊂ 1,…, 820 denote the indices of masked tokens and *ẑ***j** *the reconstructed embedding for token* ***j****. The reconstruction loss* 𝓛MAE is given by:

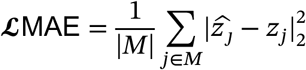

To optimize performance, we performed grid search over batch size and learning rate. The optimal configuration used a batch size of 128 and an initial learning rate of 5 × 10^−4^. A cosine annealing learning rate schedule was employed to gradually reduce the learning rate:

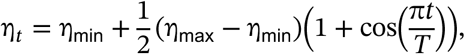

where η_*t*_ is the learning rate at step *t*, η_max_ is the initial learning rate, η_min_ is the minimum learning rate (typically set to zero), and *T* is the total number of training steps.

Training was conducted for a maximum of 200 epochs, with early stopping based on validation loss, using a patience of 20 epochs. To improve convergence stability and generalization, we applied gradient clipping with a maximum norm of 1.0 and weight decay regularization with a coefficient of 10^−2^.

The total training time was approximately 12 hours on a single NVIDIA A100 GPU, with convergence typically reached within 120 epochs.

### Hyperparameters and Model Optimization

MMT, SSM, LFADs, RSLDS ad GRU were optimized using the AdamW optimizer with a cosine annealing learning rate scheduler. A 5-fold cross-validation approach was used, selecting one mouse as the validation subject for each fold to ensure generalization. Hyperparameters were fine-tuned via grid search, exploring variations in the number of layers (2-6), learning rate (0.1 to 0.0001), dropout rates (0.1, 0.05, 0.2), and hidden dimension sizes (256, 512, 768, 1024).

Optimization algorithms compared included Adam, AdamW, and SGD. Early stopping with a patience threshold of 10 epochs was employed to prevent overfitting.

### Self-Attention Analysis

To quantify the contribution of individual cortical regions and timepoints to trial predictions, we utilized an Attention Rollout technique. This method aggregates attention weights across multiple layers of a decoder-only transformer model to derive a global representation of cortical regions interactions. By computing cumulative attention propagation, this approach enables visualization of how information is integrated across layers and which regions influence the final prediction.

Computation of Attention Rollout: Self-attention weights A^l^ were extracted from each layer ll of the decoder-only transformer model. These attention matrices had dimensions (N, H, T, T) where N corresponds to the batch size, H to the number of attention heads, and T to the sequence length. To reduce variability across heads, we computed the mean attention weight at each layer using the equation: 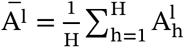 where 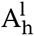 denotes the attention matrix for head h in layer l.

To obtain an effective measure of information flow through the network, attention matrices were iteratively multiplied in a top-down manner: 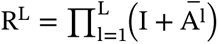 where I is the identity matrix, ensuring retention of original token contributions. The final rollout matrix was normalized such that each row summed to one, thereby preserving interpretability of token importance.

To determine the most important attention values, we employed a perturbation test by comparing trained and non-trained models. Specifically, we computed attention values for the same input using a non-trained model and established a threshold at the 95th percentile of these attention values. Attention values from the trained model that exceeded this threshold were considered significant.

The resulting rollout matrix provided a holistic view of long-range dependencies between tokens. Heatmaps were generated to visualize token-level importance, allowing for direct interpretation of model behavior. To further enhance interpretability, we focused on the attention patterns of the CLS token as it serves as an aggregate representation of the input. This visualization enabled clearer insights into the contributions of specific regions and timepoints to the final model output.

---

**Supplementary Figure 1.**
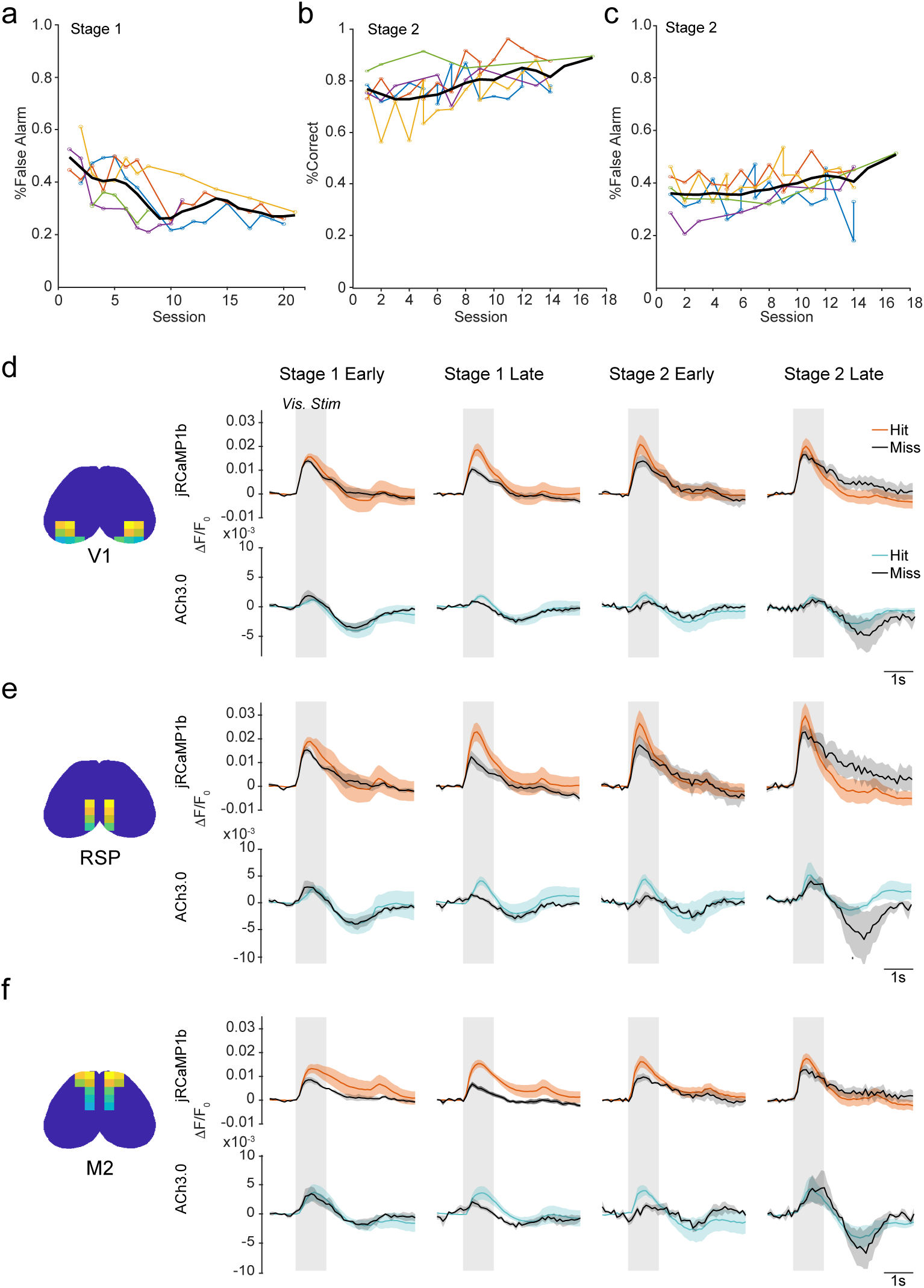
Visual task learning stages. **a.** Performance curves during Stage 1 learning. False alarm rates for each mouse are shown in different colors and the population average false alarm rate across sessions is shown as an overlaid black line. **b**. Percent correct trials in each Stage 2 training session. Percent correct rates are shown in different colors for each mouse and the population average correct rate across sessions is shown as an overlaid black line. **c**. Percent false alarm trials for each Stage 2 training session. False alarm rates for each mouse are shown in different colors and the population average false alarm rate across sessions is shown as an overlaid black line. **d**. Population average response in primary visual cortex (V1) during Early and Late sessions in Stage 1 and Stage 2 for jRCaMP1b (upper) and ACh3.0 (lower signals. In each case, signals during hit trials are denoted in color and signals during miss trials are denoted in black. **e**. Same as panel d, for activity in retrosplenial cortex (RSP). **f**. Same as panel d, for activity in secondary motor cortex (M2).

**Supplementary Figure 2.**
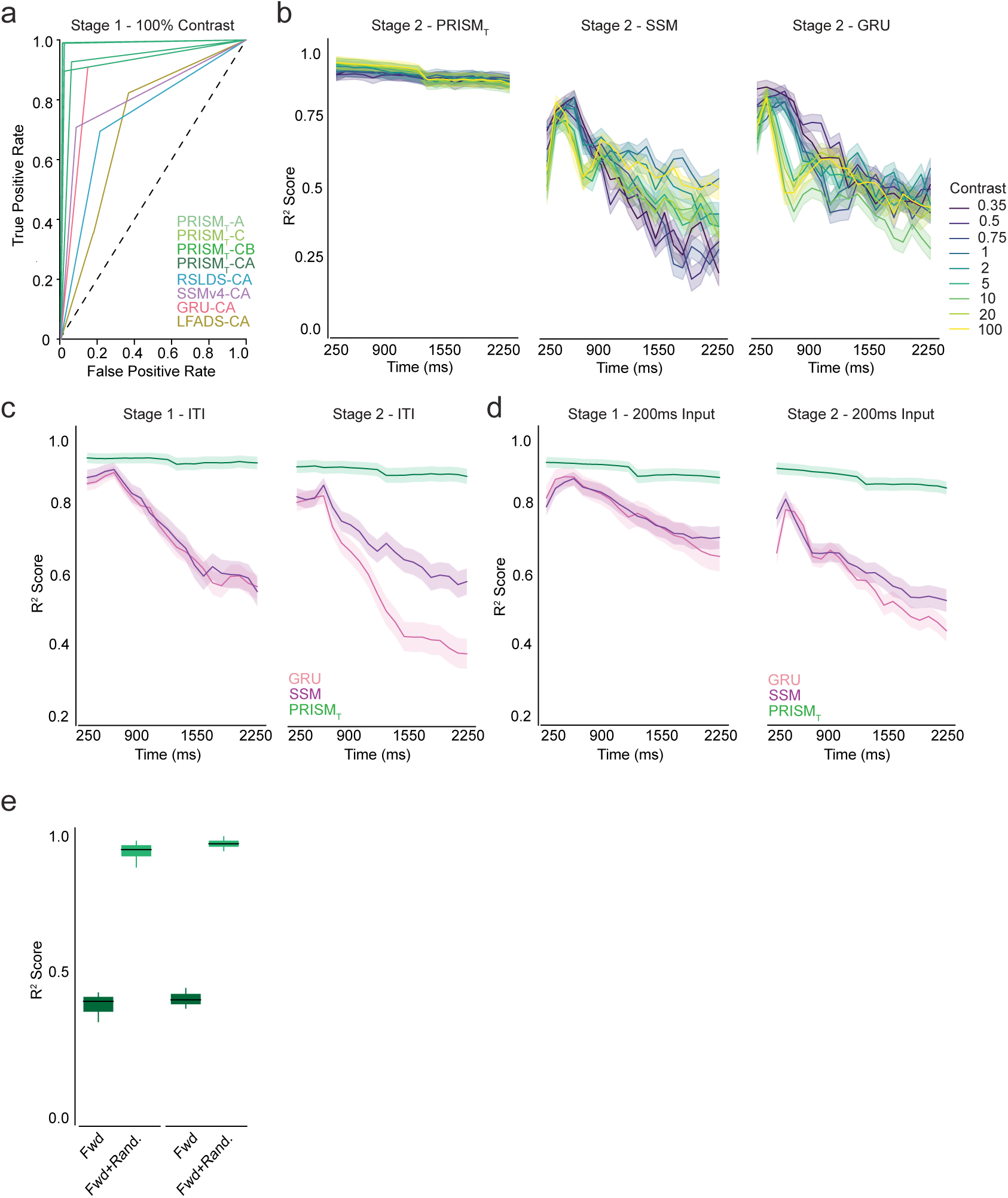
PRISM_T_ predicts neural and behavioral dynamics through learning. **a.** Stage 1 AUC-ROC curve for trial-to-trial behavior prediction. PRISM_T_ outperforms other models (n=80 recording sessions, 2911 trials, 5 mice, p < 0.05, DeLong’s test). **b.** R2 per timepoint for different contrast per model. Decrease in predictability for low contrast trials for GRU and SSM compared to PRISM_T_ (n=80 recording sessions, 3200 trials, 5 mice, p < 0.05, WSR test). **c.** Inter-trial interval forward prediction. **d.** R2 per timepoint for different models for input of 200 ms (2 frames). Performance below 400 ms model (Figure 2C), however, PRISM_T_ maintains high performance across timepoints (n=80 recording sessions, 3200 trials, 5 mice, p < 0.01, WSR test).

**Supplementary Figure 3.**
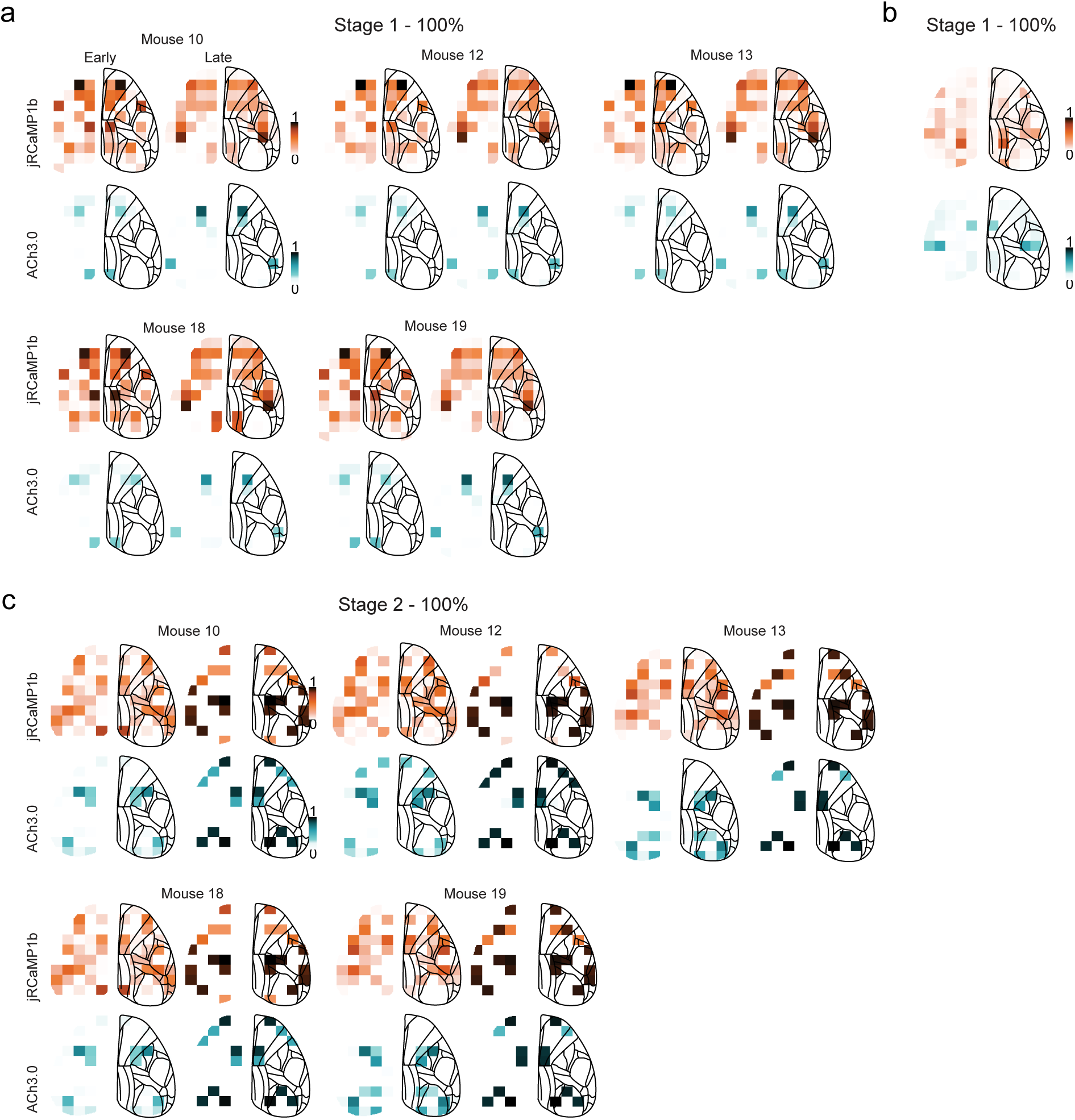
Context dependent dynamics during 100% contrast trials. **a.** Self-attention values for prediction of Hit vs Miss trials using jRCaMP1b (orange) and ACh3.0 (cyan) during stimulus presentation for late stage 1 (last 4 days) for 100% contrast trials by mice (per parcel n=32 recording sessions, 1000 trials, 5 mice, p < 0.05, permutation test). **b.** Self-attention values for predicting if a correct withholding trial, where the mouse withholds a lick when no stimulus is on the screen, comes from Early or Late Stage 1 using jRCaMP1b (orange) and ACh3.0 (cyan) during stimulus presentation for 100% contrast trials. **c.** Self-attention values for prediction of Hit vs Miss trials using jRCaMP1b (orange) and ACh3.0 (cyan) during stimulus presentation for late stage 2 (last 4 days) for 100% contrast trials by mice (per parcel n=20 recording sessions, 500 trials, 5 mice, p < 0.05, permutation test).

**Supplementary Figure 4.**
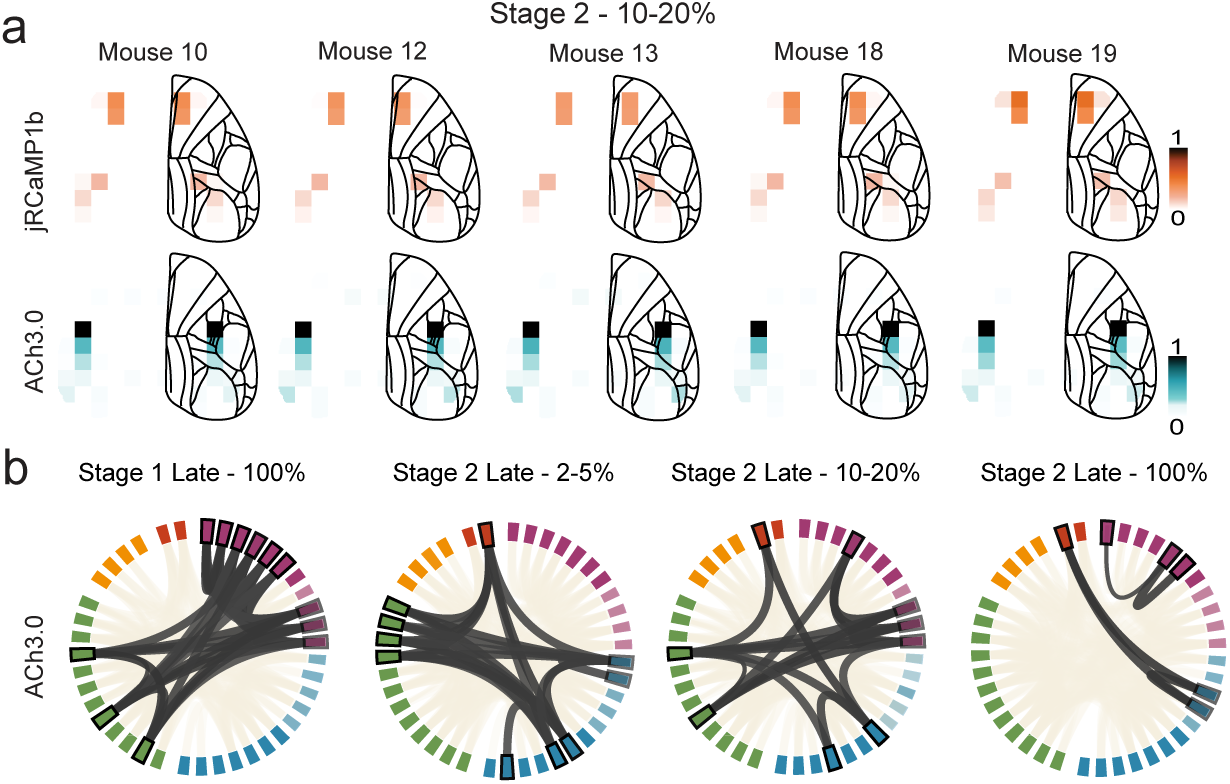
Pre stimulus onset dynamics predicts trial-to-trial mice performance. **a.** Self-attention values for prediction of Hit vs Miss trials using jRCaMP1b (orange) and ACh3.0 (cyan) using pre-stimulus dynamics for late stage 2 (last 4 days) for 10-20% contrast trials by mice. **b.** Chord plot visualizing statistically significant functional connectivity between model selected grids for Late Stage 1 (100% contrast), Late Stage 2 (2-5% contrast), Late Stage 2 (10-20% contrast), and Late Stage 2 (100% Contrast) for ACh3.0 signal. Bold indicates model selected grids, named after Allen Institute Atlas area the grid belong in. Grey indicates model-selected significant grids, and beige indicates significant non-model selected grid connections (n= 8405, 5 mice, p < 0.05, linear mixture effect model).

**Supplementary Figure 5.**
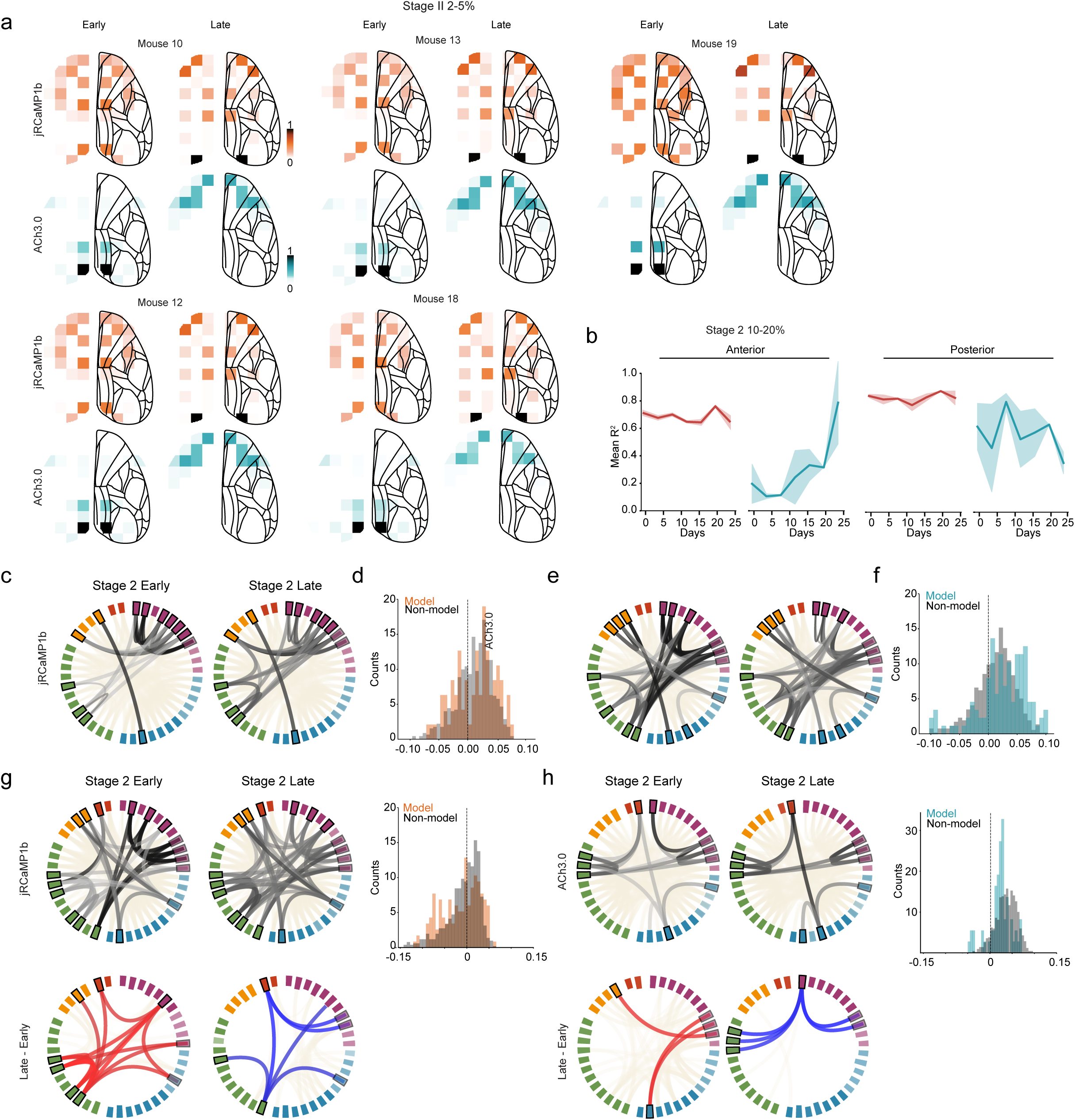
Changes in attention and dynamics associated with learning. **a.** Self-attention values for prediction of Hit vs Miss trials using jRCaMP1b (orange) and ACh3.0 (cyan) during stimulus presentation for Early Stage 2 training days (Day 1-4) for 2-5% contrast trials per mice. **b**. R2 across days for random masking during learning between anterior and posterior brain areas for 10-20% contrast trials. Split was selected based by using retrosplenial cortex as a boundary. jRCaMP1b R2 maintains predictability across days for anterior and posterior areas but ACh3.0 increases in R2 by days for anterior and decreases for posterior areas. **c.** Chord plot visualizing functional connectivity between model-selected grids for Early and Late Stage 2 for 2-5% contrast trials. Dark grey indicates model-selected grids for jRCaMP1b and ACh3.0 signals, respectively (n= 8405, 5 mice, p < 0.05, linear mixture effect model). **d.** Chord plot visualizing statistically significant positive and negative changes in functional connectivity between model selected grids (n = 8405, p < 0.05, linear mixture effects model) between Early and Late Stage 2 for 10-20% contrast trials for calcium. Bold indicates model-selected grids. Red indicates positive change, blue negative changes, and beige indicates significant non-model selected grid connections. Histogram of connectivity changes for model grids and non-model grids for calcium. No significant changes in connectivity distribution between model and non-model grids (n=800, p=0.18, linear mixture effects model). Significance was computed via Mann-Whitney U test. **e.** Chord plot visualizing statistically significant positive and negative changes in functional connectivity between model selected grids (n = 8405, p < 0.05, linear mixture effects model) between Early and Late Stage 2 for 10-20% contrast trials for ACh3.0. Bold indicates model-selected grids. Red indicates positive change, blue negative changes, and beige indicates significant non-model selected grid connections. Histogram of connectivity changes for model grids and non-model grids for jRCaMP1b. Significant changes in connectivity distribution between model and non-model grids (p << 0.001). Significance was computed via Mann-Whitney U test.

